# Reducing blood culture contamination using an initial specimen diversion device

**DOI:** 10.1101/316844

**Authors:** Frederic S. Zimmerman, Marc V. Assous, Shoshana Zevin, Yonit Wiener-Well

**Affiliations:** Department of Intensive Care, Shaare Zedek Medical Center, affiliated with the Hebrew University-Hadassah Medical School, Jerusalem, Israel; Laboratory of Clinical Microbiology and Immunology, Shaare Zedek Medical Center, affiliated with the Hebrew University-Hadassah Medical School, Jerusalem, Israel; Department of Internal Medicine B, Shaare Zedek Medical Center, affiliated with the Hebrew University-Hadassah Medical School, Jerusalem, Israel; Infectious Disease Unit, Shaare Zedek Medical Center, affiliated with the Hebrew University-Hadassah Medical School, Jerusalem, Israel

**Keywords:** blood culture, Contamination, diversion device

## Abstract

False positive blood cultures result from contamination, consuming microbiological laboratory resources and causing unnecessary antibiotic treatment and lengthened hospitalizations. Skin sterilization has been shown to reduce contamination; however, bacteria do not only colonize the surface of human skin, but are also found in deeper tissues, requiring additional techniques to reduce contamination. An initial specimen diversion device diverts the initial 1-2 ml of blood so as to remove any potentially contaminated skin plug, thus potentially further reducing culture contamination. The device has been associated with a reduction in culture contamination over short study periods in certain populations. However, more study is needed to understand whether the effect continues over longer periods of time and in hospitalized patients. Thus, in this prospective, controlled pragmatic study, cultures were obtained from hospitalized patients using the initial specimen diversion device, with cultures taken using standard methods serving as control. In total, 671 blood cultures were obtained: 207 cultures were taken using an initial specimen diversion device, with 2 (1.0%) contaminated cultures and 464 cultures were taken without the device, with 24 (5.2%) contaminated cultures (p < 0.008). No significant difference was shown in the rate of true positive cultures. Thus, use of a diversion device was associated with reduced culture contamination in hospitalized patients over a six month period without concomitant reduction in true positive cultures. This intervention may result in a reduction in costs, antibiotic use and duration of hospitalization.

## Introduction

Bloodstream infections cause significant morbidity and mortality and their prompt identification is an essential part of modern medicine. Blood cultures, first described in the latter part of the nineteenth century^1^, are an essential element of the diagnosis and treatment of patients with such infections. However, as with all medical testing, false positive results occur and these result in delays in diagnosis, inappropriate treatment and significantly added expenses.^2,3^ False positive results in blood cultures have been described for as long as there have been such cultures and are primarily due to contaminants.^1,2^ It has been estimated that up to 50% of positive blood cultures represent contamination.^2,3^ These false positive cultures, at the microbiological laboratory level, require significant additional resources for workup. In addition, and perhaps more importantly, these false positive cultures may result in unnecessary antibiotic treatment as well as additional hospitalization days, causing needless harm to patients. All this together results an estimated additional cost of $4,385-$8,720 per false positive blood culture in the United States.^3–6^

Various methods have been implemented in order to reduce blood culture contaminants. Overall, combined interventions have proven effective in attenuating blood culture contamination and many institutions have successfully reducing blood culture contamination to the currently recommended rate of less than 2-3%, with the highest achieving institutions having rates as little as 0.6%.^4,7^ However, it has been shown that the bacteria which colonize the human skin are found not only on the surface but in fact in deeper tissues as well. It has further been shown that these bacteria cannot be completely removed even under ideal conditions.^7,8^ Thus zero or near zero blood culture contamination using current methods may be impossible, even with perfect technique. As such, it is reasonable to seek techniques beyond skin sterilization and improved adherence in order to further reduce the rate of blood culture contamination.

Recently, a new device was developed that diverts the initial 1-2ml blood so as to remove any potential skin plug with contaminants from entering the blood culture bottle. The SteriPath device (Magnolia Medical Technologies, Seattle, WA) is a closed-system, sterile blood collection system that diverts 1-2mL of the initial venipuncture blood into an isolated diversion chamber and then allows venous blood to flow into culture bottles. This initial specimen diversion device has been tested in some initial studies and has been shown to reduce blood culture contamination without reducing sensitivity to true blood stream infection. A recent study showed the utility of this device in an emergency room setting.^9^ However, this study was conducted over a short period of time, did not evaluate in-patients and used paired cultures, rather than a pragmatic design. Thus this study was designed as a pragmatic trial to evaluate the use of the diversion device in reducing blood culture contamination over the period of approximately 6 months in an in-patient population.

## Methods

### Study design

This prospective, controlled, pragmatic study was conducted in the Shaare Zedek Medical Center, a 1000-bed university-affiliated general hospital in Jerusalem, Israel. The hospital includes all major departments and services, including four medical and two geriatric wards, hematology and oncology, pediatrics, a surgical division including a vascular surgery unit, gynecology and obstetrics, cardiothoracic surgery, urology, orthopedics, plastic surgery, ophthalmology, otorhinolaryngology, neurosurgery and several intensive care units. The study was conducted in the 6-month period from March 2017 through August 2017 in one of the departments of medicine (Medicine B) in our institution.

During study period, as is standard practice in our institution, blood cultures were, in preference, taken by phlebotomy teams using venipuncture, two interventions that have been shown in multiple studies to reduce the rate of blood culture contamination.^2,6^ The remainder of cultures were taken mostly by resident physicians. Prior to study initiation a feedback mechanism was initiated in our institution in order to reduce contamination prior to the study (unpublished data) and reduce observer effect during the study period.

During the course of the study, in a convenience sample of patients in one medical department, blood cultures were obtained using the initial specimen diversion device, either via an integrated needle, or in cases of intravenous line placement, by attaching the device to the newly placed intravenous catheter. The control group consisted of blood cultures taken in the department using standard devices – Vacuette Holdex^®^ (Grenier Bio-One, Kremsmünster, Austria) attached to either a scalp vein set or to an intravenous catheter. In either method, blood was inoculated into BD BACTEC™ Plus Aerobic/F in addition to Plus Anaerobic/F culture bottles (BACTEC, Franklin Lakes, NJ). An automated blood culture system, BD BACTEC FX (BACTEC) was used to process the blood cultures. Relevant characteristics of the blood culture, including date of culture and site of blood collection, were recorded. Isolates were identified using MALDI-TOF (Vitek MS, bioMérieux) and/or standard biochemical methods.

Blood culture contamination was identified by microbiological criteria – growth of coagulase-negative staphylococci (CNS), corynebacteria, micrococci or alpha-hemolytic streptococci. Cultures could be reclassified as true infections if such an organism was isolated from multiple blood cultures obtained by different venipuncture. It should be noted that given the high sensitivity and short time to positivity of modern blood culture systems ^10^, time to positivity is not an effective method of identifying culture contaminants. If a single blood culture showed growth of true infection and contaminant organisms, it was included in both categories.

During the first month of the study observers provided technical support to ensure proper use of the device by the users – whether phlebotomists or physicians. The observers did not provide direct assistance in blood culture collection. For the remaining five months of the study the device was used without observer presence. During the entire period, blood cultures were collected according to the clinical discretion of the treating physician.

Ease of use of the device was evaluated using a standardized survey of phlebotomists and others involved in sample collection.

Data was analyzed using Win-Pepi (version 11.65). Proportions were compared using the χ^2^-score or Fisher’s exact test, where appropriate. Continuous variables were compared by Student’s t-test or Mann-Whitney-Wilcoxon test where appropriate. Two-tailed p values were taken and a p value <0.05 was defined as significant.

The study was approved by the Institutional Review Board of Shaare Zedek Medical Center.

## Results

In the 6 months of the study period, 671 blood cultures were obtained in the study department. Of these, 207 cultures were recorded to have been taken using the initial specimen diversion device; this population served as the study group. In 464 cultures, use of the device was not recorded; this population served as the control group. A significant difference in age was found between the control group and the study group (74±1.4 versus 77±2.1, respectively, p<0.004). Otherwise no differences between the groups were noted (Table 1).

**Table 1.** Characteristics of study population. CI: confidence interval

| **Variable** | **Diversion device**<br>**n=207 (%)** | **Standard practice**<br>**n=506 (%)** | **p-**<br>**value** |
| --- | --- | --- | --- |
| Male | 118 (57.0) | 311 (61.5) | 0.446 |
| Age - years±95% CI | 77±2.1 | 74±1.4 | 0.004 |
| Place of residence |  |  |  |
| home | 148 (71.5) | 385 (76.1) | 0.292 |
| assisted living | 11 (5.3) | 23 (4.5) | 0.847 |
| chronic care facility | 48 (23.2) | 98 (19.4) | 0.303 |
| Hospitalized in last year | 128 (61.8) | 304 (60.1) | 0.669 |
| Seven day mortality | 36 (17.4) | 77 (15.2) | 0.498 |

Of the 464 cultures in the control group, 68 (14.7%) were positive for bacterial growth. Of those positive cultures, 24 were defined as contaminants, thus resulting in a contamination rate of 5.2%. In the study group, 207 blood cultures were obtained. Of these, 18 (8.7%) were positive for bacterial growth. Of these positive cultures, 2 were defined as contaminants, thus resulting in a contamination rate of 1.0% in the study group (p<0.008 for the difference between the groups). With the exception of a single culture in the control group which grew *Streptococcus parasanguinis*, all other contaminated cultures grew coagulase-negative Staphylococci.

Of the control group cultures, 44 were defined as true positives resulting in a true positive rate of 9.5%. Of the study group cultures, 16 were defined as true positives resulting in a true positive rate of 7.7%. Enterobacteriaceae and staphylococci were the most common causes of true bacteremia in both groups. No significant difference in the true positive rate or in the microbiological characteristics of true positive cultures were noted between the groups (Table 2 and Figure 1).

**Figure 1.**
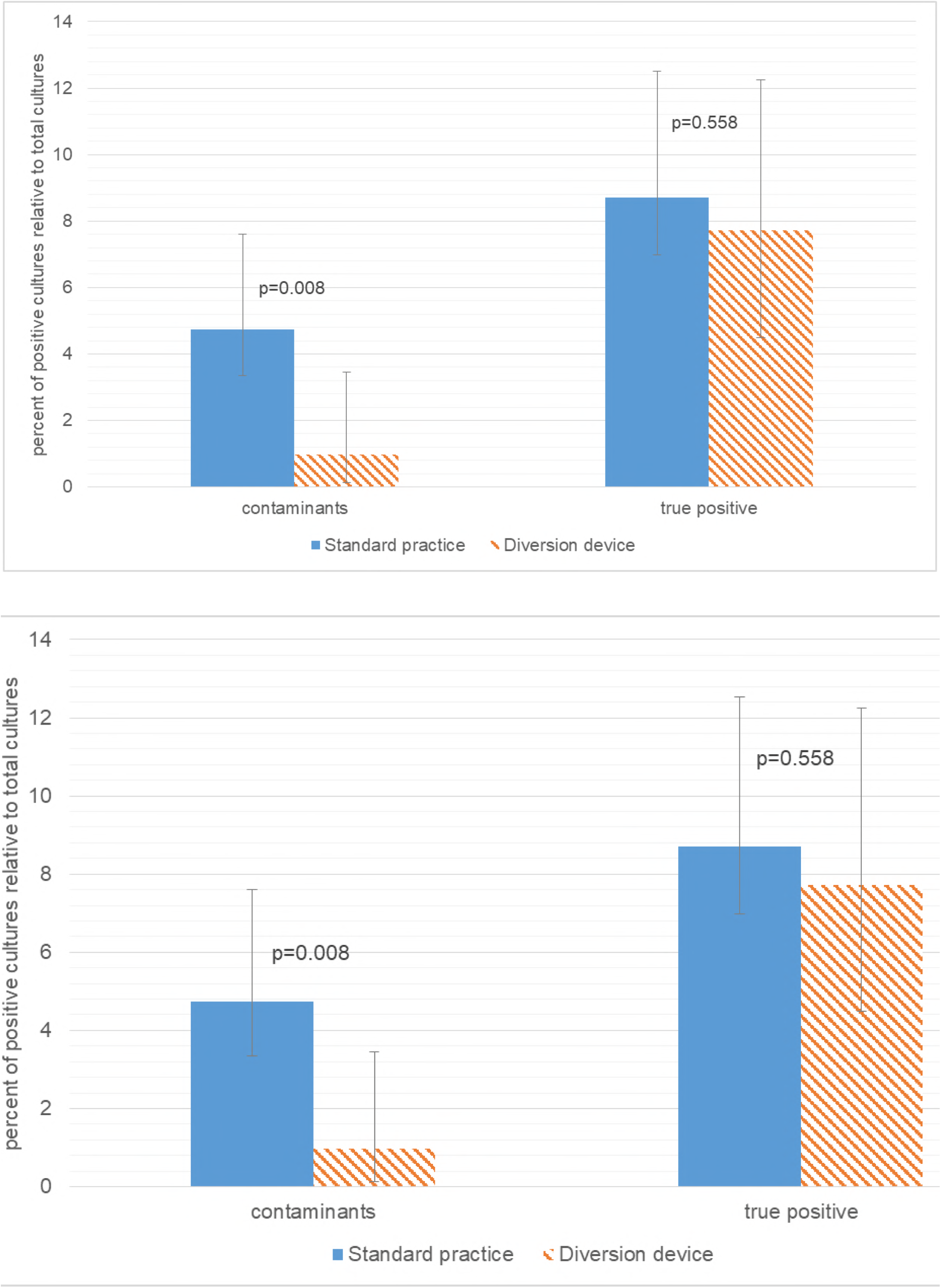
Positive culture rate for both contaminated and true positive cultures.

**Table 2.** Microbiological and clinical characteristics of positive cultures. (CoNS: coagulase negative staphylococci)

| **Variable** | **Diversion device**<br>**n=207 (%)** | **Standard practice**<br>**n=464 (%)** | **p-value** |
| --- | --- | --- | --- |
| All positive cultures | 18 (8.7) | 68 (13.4) | 0.034 |
| Culture contamination | 2 (1.0) | 24 (4.7) | 0.008 |
| CoNS | 2 (1.0) | 23 (4.5) | 0.013 |
| Other | 0 (0.0) | 1 (0.2) | 1.000 |
| Contamination by age group |  |  |  |
| 18-64 | 0/28 (0.0) | 2/96 (2.1) | 1.000 |
| 65-80 | 1/70 (1.4) | 16/226 (7.1) | 0.081 |
| 81+ | 1/109 (0.9) | 6/184 (3.3) | 0.253 |
| True positive | 16 (7.7) | 44 (8.7) | 0.558 |
| Enterobacteriaceae | 3 (1.4) | 20 (4.0) | 0.067 |
| Staphylococci | 12 (5.8) | 18 (3.6) | 0.312 |
| Other | 1 (0.5) | 6 (1.2) | 0.447 |

In a survey of ease of device use, all respondents who had used the device more than once considered it slightly or moderately more difficult to use than standard practice (grade 2 or 3 out of 5).

## Discussion

A substantial fraction of positive blood cultures represent contamination, rather than true bloodstream infection^2,3^, including in the control group of our study (Figure 1), where 24 out of 464 (5.2%) cultures represented contamination and 44 (9.5%) represented true infection. These false positive cultures, at the microbiological laboratory level, require significant futile resources for workup. In addition, and perhaps more importantly, these false positive cultures result in unnecessary antibiotic treatment as well as excessive hospitalization days, causing needless harm to patients. All this together results in substantial additional costs per false positive blood culture.^3–6^

In this study, we investigated the efficacy of an initial specimen diversion device in reducing the rate of blood culture contamination. We found that the device was associated with a reduction in blood culture contamination with a 5.2% (24 out of 464 cultures) contamination rate in the control group and a 1.0% (2 out of 207 cultures) contamination in the study group (p<0.008). These results are consistent with those obtained in a recent study performed in an emergency room population.^9^ In our study, the control group was noted to be younger than the study group.

This study suggests that use of an initial specimen diversion device can effectively reduce culture contamination among hospitalized patients without impacting the rate of true positive cultures. Furthermore, this reduction can be maintained over a relatively extended period of time.

The study has several limitations. First, though this study was controlled, a pragmatic design using a convenience sample, rather than a randomized control, was employed. Such a study is limited in controlling for confounders related to selection of sample. Indeed, the study population was older than that of the control, though no other significant differences were noted (Table 1). Additionally, this study was a single center study with a relatively small study population. As such, results should be interpreted cautiously.

In conclusion, this study found that use of an initial specimen diversion device was associated with a reduction in blood culture contamination in hospitalized patients over a six month period – from 5.2% (24/464) in the control group to 1.0% (2/207 – p<0.008) in the study group – without a concomitant reduction in true positive cultures. This is an intervention that can result in a reduction in costs, unnecessary antibiotic use and duration of hospitalization.

## Acknowledgments

The authors would like to thank Dr. David Raveh for assistance in statistical analysis.

The funding source, Magnolia Medical Technologies, provided the SteriPath device as well as initial technical support in their use. The funding source did not provide any other financial support nor were they involved in study design; the collection, analysis and interpretation of data; in the writing of the report; or in the decision to submit the article for publication.

